# A computational framework to assess genome-wide distribution of polymorphic human endogenous retrovirus-K in human populations

**DOI:** 10.1101/444034

**Authors:** Weiling Li, Lin Lin, Raunaq Malhotra, Lei Yang, Raj Acharya, Mary Poss

## Abstract

Human Endogenous Retrovirus type K (HERV-K) is the only HERV known to be insertionally polymorphic. It is possible that HERV-Ks contribute to human disease because people differ in both number and genomic location of these retroviruses. Indeed viral transcripts, proteins, and antibody against HERV-K are detected in cancers, auto-immune, and neurodegenerative diseases. However, attempts to link a polymorphic HERV-K with any disease have been frustrated in part because population frequency of HERV-K provirus at each site is lacking and it is challenging to identify closely related elements such as HERV-K from short read sequence data. We present an integrated and computationally robust approach that uses whole genome short read data to determine the occupation status at all sites reported to contain a HERV-K provirus. Our method estimates the proportion of fixed length genomic sequence (*k-mers*) from whole genome sequence data matching a reference set of *k-mers* unique to each HERV-K loci and applies mixture model-based clustering to account for low depth sequence data. Our analysis of 1000 Genomes Project Data (KGP) reveals numerous differences among the five KGP super-populations in the frequency of individual and co-occurring HERV-K proviruses; we provide a visualization tool to easily depict the prevalence of any combination of HERV-K among KGP populations. Further, the genome burden of polymorphic HERV-K is variable in humans, with East Asian (EAS) individuals having the fewest integration sites. Our study identifies population-specific sequence variation for several HERV-K proviruses. We expect these resources will advance research on HERV-K contributions to human diseases.

**Author summary:** Human Endogenous Retrovirus type K (HERV-K) is the youngest of retrovirus families in the human genome and is the only group that is polymorphic; a HERV-K can be present in one individual but absent from others. HERV-Ks could contribute to disease risk but establishing a link of a polymorphic HERV-K to a specific disease has been difficult. We develop an easy to use method that reveals the considerable variation existing among global populations in the frequency of individual and co-occurring polymorphic HERV-K, and in the total number of HERV-K that any individual has in their genome. Our study provides a global reference set of HERV-K genomic diversity and tools needed to determine the genomic landscape of HERV-K in any patient population.

## Introduction

Endogenous retroviruses (ERVs) are derived from infectious retroviruses that integrated into a host germ cell at some time in the evolutionary history of a species [1–5]. ERVs in humans (HERVs) comprise up to 8% of the genome and have contributed important functions to their host [6–8]. The infection events that resulted in the contemporary profile of HERVs occurred prior to emergence of modern humans so most HERVs are fixed in human populations and those of closely related primates. However some HERVs retain the ability to replicate and reintegrate into germline so that individuals differ in the number and genomic location occupied by an ERV, a situation termed insertional polymorphism [9–11]. Among all families of HERVs, HERV-K is the only one known to be insertionally polymorphic in humans.

A full-length retroviral sequence is called a provirus and encodes several viral structural or regulatory proteins that are flanked by two long terminal repeats (5’ or 3’ LTR). While there are several HERV-K that are full length, none are infectious and most contain mutations or deletions that affect the open reading frames or truncate the virus. Further, the identical LTRs are substrates for homologous recombination, which deletes virus genes while retaining a single, or solo, LTR at the integration site [12–14]. Thus, in a population, a site could be unoccupied, occupied by a HERV-K provirus, or contain a solo LTR. Insertional polymorphism typically refers to the occupancy at a loci [15,16]. However the occupied site can contain a provirus or solo LTR and a provirus sequence can vary among individuals. Thus HERV-K and other HERVs can contribute to genomic diversity in the global human population in several ways [17].

HERVs from multiple families have been linked with both proliferative and degenerative diseases in humans [18–24]. Although there are known mechanisms by which a HERV can cause disease_;_ for example, by inducing genome structural variation through recombination [25–29], affecting host gene expression [30], and inappropriate activation of an immune response by viral RNA or proteins [21], it has been difficult to establish an etiological role of a HERV in any disease. HERV-K specifically has been associated with breast and other cancers [3, 31–35], and autoimmune diseases, such as rheumatoid arthritis [36,37], multiple sclerosis [20,38] and systemic lupus erythematosus [8,20,39] without definitive evidence of causality or of the specific loci involved. Recently, a HERV-K envelope protein was shown to recapitulate the clinical and histological lesions characterizing Amyotropic Lateral Sclerosis [40,41], providing an important mechanistic advance of a role for a HERV-K protein in a disease.

In this paper, we focus on characterizing the genome landscape of known insertionally polymorphic HERV-K proviruses in the 1000 Genomes Project (KGP) data. We present a data-mining tool and a statistical framework that accommodates low depth data characteristic of the KGP - and often patient - data to estimate the presence or absence of a provirus at known HERV-K loci. Because combinations of HERV-K may act synergistically in the pathogenesis of a disease [42], we estimate the co-occurrence of polymorphic HERV-K proviruses in different populations and provide a tool to visualize HERV-K co-occurrence in global populations. Our results provide a reference of global population diversity in HERV-K proviruses at all currently known loci in the human genome and demonstrate that there are notable differences among population frequencies of HERV-Ks and the total number of HERV-Ks found in a person’s genome.

## Results

### A model to estimate polymorphic HERV-K from whole genome sequence data

The goal of this research was to develop a computationally efficient and easy to use tool that could accurately report the status of all HERV-Ks with coding potential (provirus) from whole genome sequence (WGS) data. We use the KGP database to establish the global population diversity of each polymorphic HERV-K and the burden of HERV-K in individual genomes to provide a foundation to study the role of HERV-K in human disease. Our method takes as input all reads that map to identified HERV-K elements in hg19. The rationale here is that polymorphic HERV-K are very similar to those in the reference genome and will map on existing elements. The recovered reads are reduced to *k-mers* and mapped to a reference set of *k-mers* representing all unique sites in every HERV-K in the database. The output is a ratio of subject *k-mers* (n) that are 100% match to the reference *k-mers* (T) (see methods for full details).

Our preliminary analysis of the KGP data demonstrated that our *k-mer*-based approach is sensitive to sequence depth; some HERV-K are represented by an almost continuous range of n/T from 0-1 (Fig 1A), making presence/absence classification difficult. A comparison of a subset of the 28 individuals in the KGP data that have both low and high sequence depth data shows how depth affects n/T (Fig 1B, see S2 Fig for data of all 28 persons). If read depth is greater than 20, there is less dispersion of n/T values, most likely because more reads are recovered from the mapped intervals. However, the majority of the KGP data is approximately 6x depth and thus to make use of this important resource, we developed a mixture model to cluster the n/T values from genomes sequenced at low depth. K was optimized to 50 because this value improved our model computational efficiency and output (Fig 1B, S1-S3 Methods, S1 Fig). The states, ‘provirus’, ‘solo LTR’, and ‘absent’ are preliminarily assigned to each cluster based on the high depth data (Fig 1B). Individuals with n/T=1 have the reference allele and n/T=0 indicates that the HERV-K is absent (no *k-mer*s to unique sites in the HERV-K were recovered from mapped sequence reads). The *k-mer*s derived from persons with low and intermediate n/T values were mapped to each HERV-K to determine whether they localized only in the LTR (assign ‘solo LTR’) or in the coding region (assign ‘provirus’) (S3 Fig).

**Fig 1.**
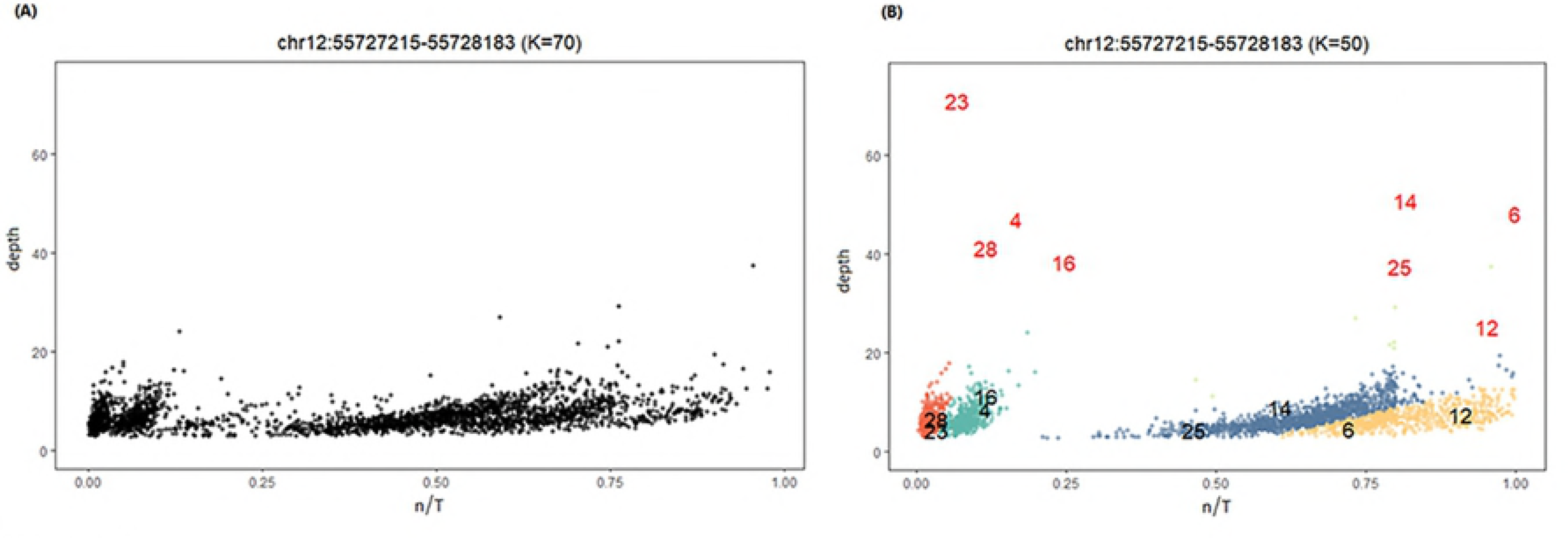
A mixture model to account for low depth WGS data. A) The plot displays the n/T value for 2535 individuals from KGP with low depth sequence data for chr12:55727215-55728183 when K=70. There is little resolution of values to enable assignment to provirus, solo LTR, or absent states. B) The result of the mixture model on the same data. The individual clusters generated by the model are indicated by a unique color; in the example shown, there are four clusters. K has been optimized to 50 to enable clear clustering in the ‘absent’, ‘solo LTR’ and ‘provirus’ states. In this example, eight of the 28 individuals that have both low and high depth sequence data (see S1_Dataset:KGP) are shown to demonstrate the effect of sequence depth. The n/T ratio is 1 for persons with high depth data [red numbers, #6 and 12] who have the reference allele, while the corresponding low depth data [black numbers, yellow cluster] from the same individuals have n/T ranging from 0.7 to 0.9. There is less of an effect of depth for individuals who do not have the HERV-K (n/T=0). However, optimizing K facilitates separation of clusters for absent [red cluster, #23 and 28], and solo LTR [green cluster, #4 and 16]. States are confirmed by mapping the *k-mer*s from individuals in a cluster to the reference HERV-K (S3 Fig).

### Prevalence of polymorphic HERV-K in each KGP super-population

The WGS data of each individual in the KGP dataset were evaluated using our analysis workflow. HERV-Ks on chrY were not considered. Twenty sites, omitting one at chr1:73594980 [see methods] were identified that were polymorphic for containing a HERV-K provirus. A phylogenetic analysis of all HERV-Ks greater than 6 kbp shows that polymorphic HERV-Ks are closely related (S4 Fig). The prevalence of the 20 polymorphic HERV-Ks varied from 0.9% to 99.5% when averaged across the entire KGP dataset (Table 1). However, there were notable differences in prevalence at each site among the five super-populations (AFR, EAS, AMR, EUR, SAS). Of the 20, the prevalence of seven polymorphic HERV-Ks was greater than 90% and the difference between populations with the lowest and highest prevalence was less than 6.5% (Table 1). There was 100% occupancy for six of the seven high prevalence polymorphic HERV-Ks (98.8% for the seventh), indicating that the rate of conversion to solo LTR is low for viruses at these sites (S1 Table). Two polymorphic HERV-Ks had an overall prevalence of less than 10% in any population (Table 1) and we found no evidence of a solo LTR at these sites; both are found in individuals from AFR. Nine of the remaining 11 HERV-Ks are of interest because the difference between super-populations with the highest and lowest prevalence is between 28 and 80 percentage points (Table 1). Of note, the prevalence is lowest in EAS populations for the three HERV-Ks with the largest difference among super-populations.

**Table 1.** Provirus frequencies of polymorphic HERV-K.

|  | **KGP**<br>**pro-**<br>**virus**<br>**(%)** | **AFR** | **AMR** | **EAS** | **EUR** | **SAS** | **max-**<br>**min** |
| --- | --- | --- | --- | --- | --- | --- | --- |
| **<u>chr1:75842771<sup>c</sup></u>** | 42.88 | 26.76 | 56.53 | 6.02 | 68.91 | 66.80 | 62.89 |
| **chr3:112743479<sup>a</sup>** | 98.46 | 96.71 | 99.72 | 99.81 | 99.60 | 97.37 | 3.09 |
| **chr3:148281477** | 41.89 | 38.86 | 42.61 | 45.05 | 46.53 | 37.45 | 9.09 |
| **<u>chr3:185280336<sup>a</sup></u>** | 99.49 | 98.06 | 100.00 | 100.00 | 100.00 | 100.00 | 1.94 |
| **<u>chr4:69463709<sup>c</sup></u>** | 72.50 | 93.87 | 88.92 | 31.07 | 85.35 | 61.94 | 62.80 |
| **chr5:156084717<sup>a</sup>** | 99.41 | 98.36 | 99.72 | 100.00 | 99.80 | 99.60 | 1.64 |
| **chr6:57623896<sup>a</sup>** | 93.65 | 90.73 | 97.16 | 90.87 | 97.23 | 94.33 | 6.50 |
| **chr6:78427019<sup>a</sup>** | 97.71 | 95.52 | 97.16 | 99.61 | 97.23 | 99.60 | 4.10 |
| **chr7:4622057<sup>\*c</sup>** | 47.50 | 61.14 | 30.11 | 58.25 | 36.44 | 41.50 | 31.02 |
| **chr8:12316492<sup>c</sup>** | 14.08 | 32.88 | 12.22 | 0 | 15.64 | 3.04 | 32.88 |
| **<u>chr8:7355397<sup>c</sup></u>** | 18.66 | 39.16 | 12.50 | 6.02 | 11.29 | 15.99 | 33.14 |
| **<u>chr10:27182399<sup>a</sup></u>** | 99.13 | 97.46 | 99.43 | 99.81 | 99.80 | 99.80 | 2.35 |
| **chr11:101565794<sup>c</sup>** | 63.04 | 80.87 | 77.27 | 6.99 | 86.53 | 63.16 | 79.54 |
| **<u>chr12:55727215</u>** | 72.19 | 72.80 | 80.40 | 63.30 | 80.99 | 65.79 | 17.69 |
| **chr12:58721242<sup>c</sup>** | 70.73 | 58.89 | 78.41 | 60.00 | 87.33 | 75.51 | 28.43 |
| **<u>chr19:21841536<sup>c</sup></u>** | 26.98 | 39.16 | 11.93 | 32.23 | 10.69 | 32.39 | 28.47 |
| **<u>chr19:22414379<sup>c</sup></u>** | 67.77 | 89.24 | 60.80 | 56.89 | 55.84 | 67.21 | 33.40 |
| **<u>chr19:22457244<sup>b</sup></u>** | 0.87 | 3.29 | 0.00 | 0.00 | 0.00 | 0.00 | 3.29 |
| **chr22:18926187<sup>a</sup>** | 99.49 | 98.36 | 99.72 | 100.00 | 99.80 | 100.00 | 1.64 |
| **<u>chrX:93606603<sup>b</sup></u>** | 2.25 | 7.32 | 2.27 | 0.00 | 0.00 | 0.00 | 7.32 |

For simplicity, only the starting coordinate is listed.
\* The value given represents those with the tandem repeat
<sup>a</sup>: prevalence > 90%
<sup>b</sup>: low prevalence and no solo LTR
<sup>c</sup>: max-min difference is > 28%

Individuals from African populations differ significantly from the other four super-populations in the prevalence of ten of the polymorphic HERV-K, three of which occur in close proximity on chr19. (Table 1, S2_Dataset:compare_prevalence). EUR and AFR super-populations are significantly different at all but one of the 20 polymorphic HERV-K based on adjusted p-values (S2_Dataset:compare_prevalence).

### The number of polymorphic HERV-Ks per individual

The HERV-K genome is close to 10 kbp. As there are 20 HERV-Ks that are polymorphic in human populations, we asked if some individuals carry a different burden of these repetitive, and potentially functional, viral elements than others. This was indeed the case. The number of polymorphic HERV-K proviruses per person ranges from 7-18 (Fig 2, S2_Dataset:HERV-K per person). More than 63% of individuals from all super-populations except EAS carry 12 to 14 proviruses in their genome. Individuals from EAS have a lower burden with 69% of individuals carrying 9-11 HERV-K proviruses. 7% of AFR individuals have 16 or 17 proviruses compared to a maximum of 2% in other groups (S2_Dataset:HERV-K per person). These data highlight the importance of using a comprehensive approach to study the potential disease impact of polymorphic HERV-Ks, because total HERV-K burden in individuals might influence a disease phenotype without significant variation in prevalence of each provirus in a patient pool.

**Fig 2.**
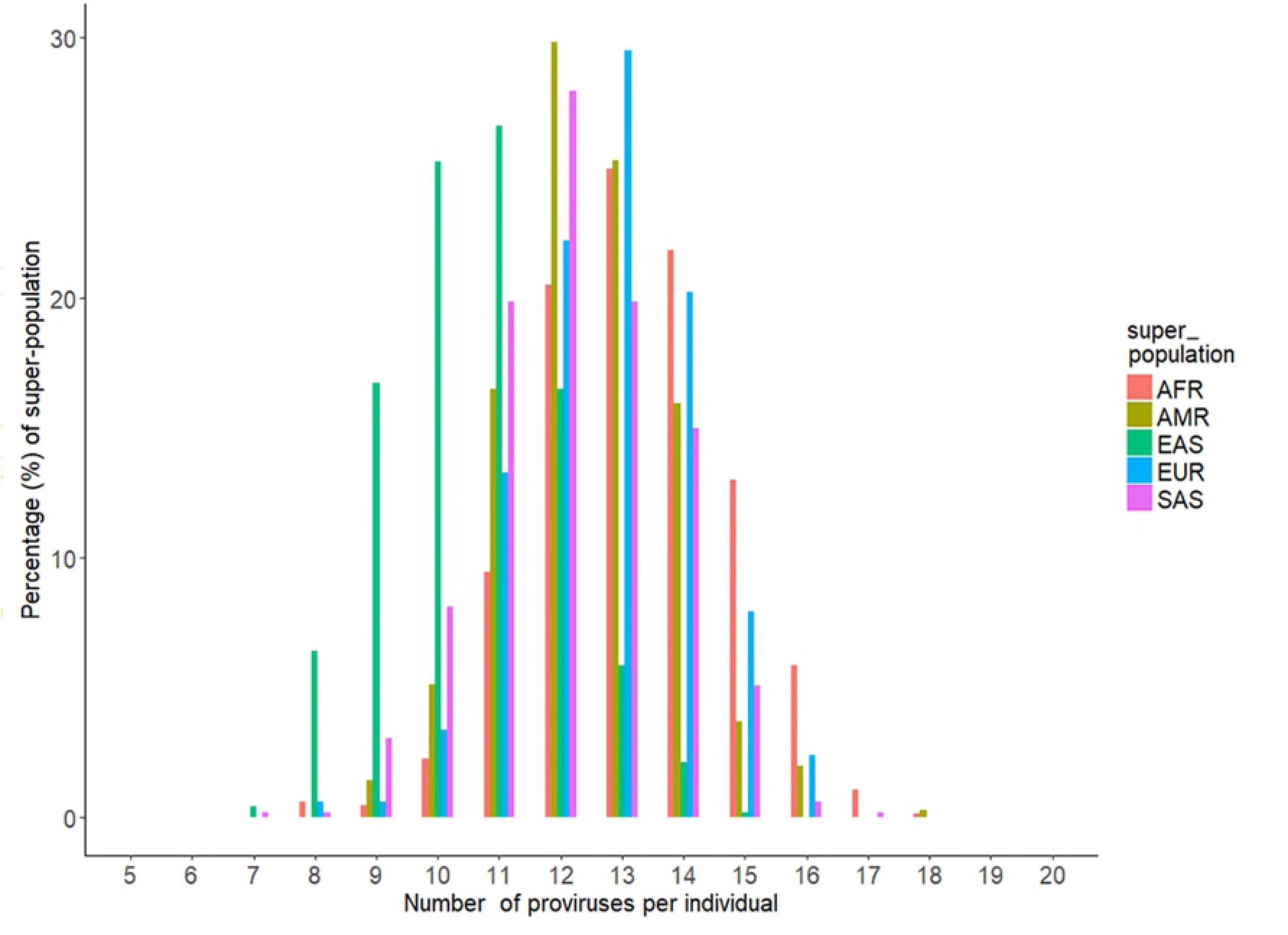
Histogram of the number of proviruses per individual among super-populations.

### Co-occurrence of polymorphic HERV-Ks

Our data provide a comprehensive picture of sites occupied by HERV-K provirus in each genome, which enables analysis of polymorphic HERV-K co-occurrence in populations. We assessed combinations of three, four and five polymorphic HERV-Ks and found that there are many combinations of co-occurring viruses that are population-specific (S3_Dataset). To facilitate exploration of HERV-K combinations among KGP populations, we developed a D3.j visualization tool that allows a user to choose any combination of the 20 polymorphic HERV-K proviruses and display the co-occurrence prevalence among the 26 populations represented in the KGP data. As an example, we present a combination of four HERV-Ks to represent the variation that occurs in KGP individuals, which in this case ranges from 3% in EAS to 59% in EUR (Fig 3A). We also determine that the three polymorphic HERV-Ks found on chr19 co-occur only from three AFR populations and in less than 2% of individuals (Fig 3B).

**Fig 3.**
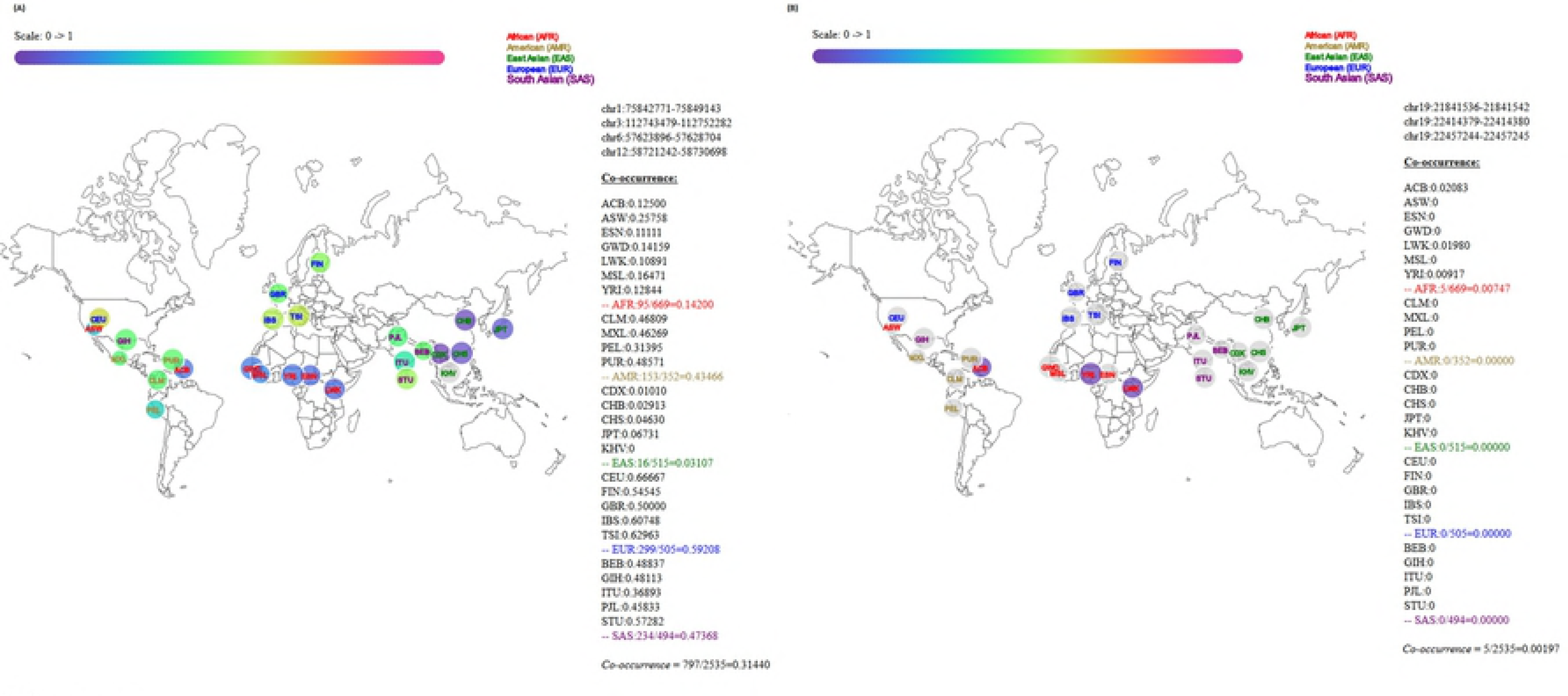
A visualization tool to examine co-occurrence of polymorphic HERV-Ks. A) The co-occurrence of HERV-Ks at chr1:75842771-75849143, chr3:112743479-112752282, chr6:57623896-57628704, and chr12:58721242-58730698 in the 26 populations are represented based on their geographic location. The relative frequency for these four co-occurring HERV-Ks in each population bubble is displayed based on the color gradient shown in the scale at the top. The actual prevalence of the given combination of HERV-K provirus for each population and the cumulative prevalence for the super-population are shown in text on the right. Note that AFR and EAS have the lowest prevalence of these four polymorphic HERV-Ks. B) As in (A) showing the co-occurrence of the three polymorphic HERV-Ks that are present on chr19 by population. This is a rare combination only found in two AFR populations and individuals in the Caribbean of African ancestry.

### HERV-K status informs KGP super-populations

Because there are clearly population-specific differences in both individual HERV-K frequency and in the frequency of HERV-K co-occurrence, we explored whether the presence or absence of polymorphic HERV-Ks is sufficient to distinguish populations using Fisher’s linear discriminant analysis (LDA) [43]. Based on the status ‘provirus’, ‘solo LTR’, or ‘absence’, there is little resolution of AFR, EUR, and EAS super-populations (Fig 4A). However, there is sufficient signature to separate AFR, EUR, and EAS if we utilize the n/T ratio on the 20 polymorphic HERV-Ks (S5 Fig) and we further improve population separation if we use the n/T ratio for all 96 HERV-Ks (Fig 4B). This indicates that we are losing information by reducing the data to three states and that fixed HERV-K also contain signal for population of origin.

**Fig 4.**
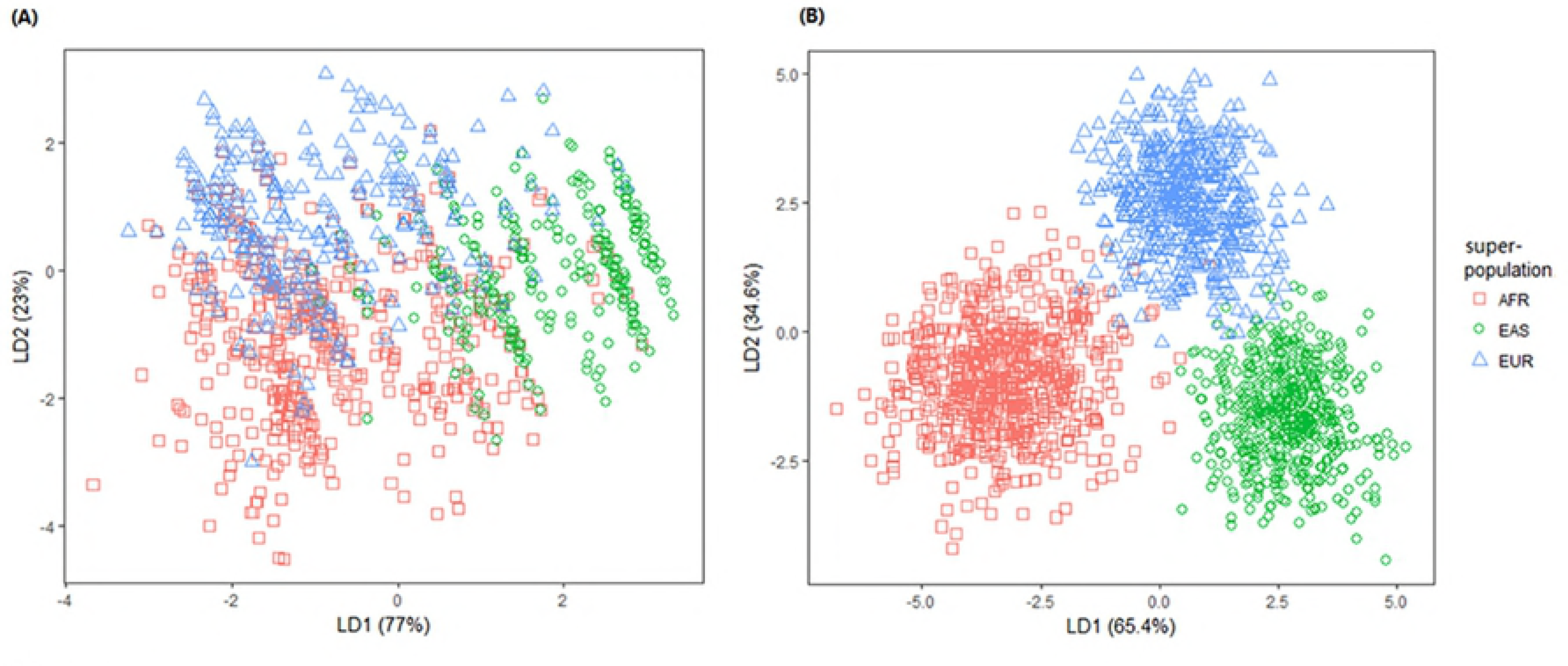
Linear discriminant analysis of HERV-K status among three super-populations. A) LDA based on the states ‘provirus’, ‘solo LTR’ and ‘absence’ of the 20 polymorphic HERV-K for AFR, EAS, and EUR. AMR and SAS overlap these three populations and are removed for clarity B) LDA plot on n/T ratio of all 96 HERV-K discriminates AFR, EAS, and EUR super-populations. See S6 Fig for plots with all five super-populations.

An n/T = 1 indicates that we recovered all reads that map to the reference set T for a specific HERV-K. If there is a HERV-K allele that has not been reported in any database but that is common in a population, we expect n/T <1 because we require 100% match to reference set T and *k-mers* covering allelic sites will not be included. We assessed the density distributions of n/T plots for each of the 96 HERV-Ks for evidence of population-specific alleles (S4 Method, S7 Fig). Five HERV-Ks have some indication of population specific distributions (S1_Dataset:virus). The HERV-K at chr1:155596457-155605636, which we report as fixed, is notable because the reference allele (n/T=1) is only found in AFR (Fig 5A, S7 Fig). Individuals from most other populations have n/T near 0.5. We mapped *k-mer*s from individuals with n/T near 0.5 to the reference HERV-K sequence and confirmed that there is a loss of *k-mer*s at several sites covered by the unique reference *k-mer*s for this virus (S8 Fig). There are also cases where the reference allele is found in all populations except AFR (Fig 5B and see S7 Fig for additional examples).

**Fig 5.**
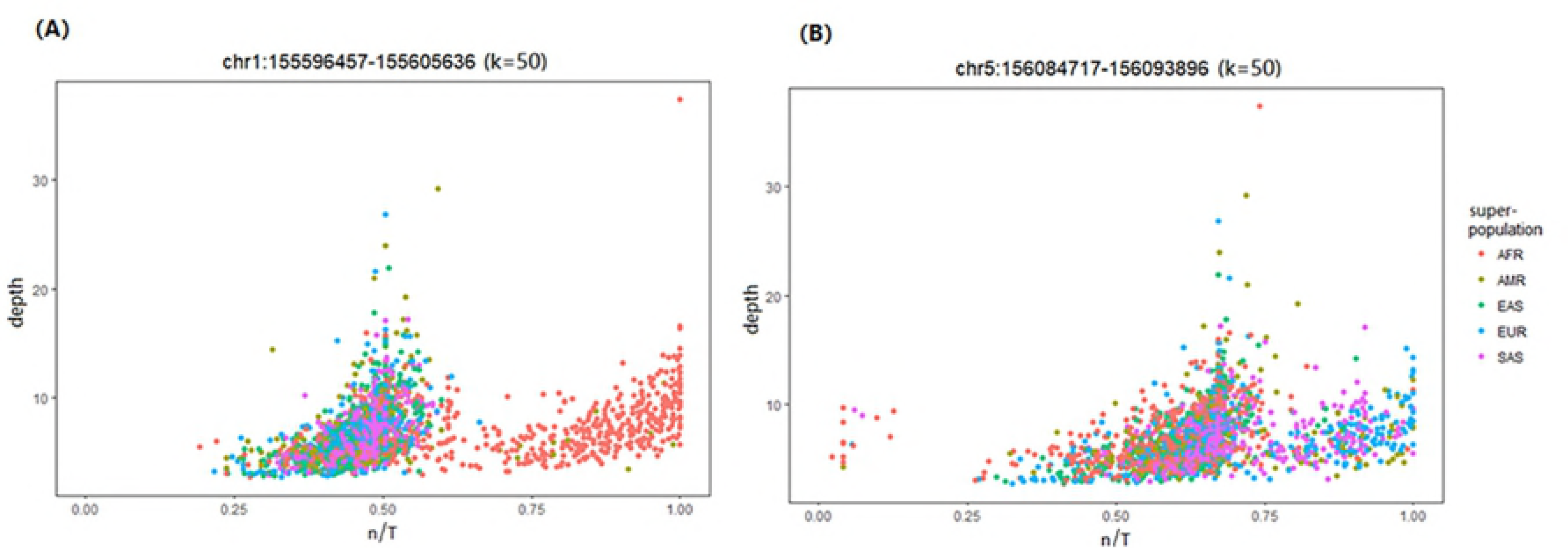
Population specificity of HERV-K alleles. A) n/T plot for chr1:155596457-155605636 colored by each of the 5 super-populations. Only individuals from AFR and a few from AMR have the HERV-K reference sequence. B) Plot of chr5:156084717-156093896 colored by each of the 5 super-populations. In this case, all populations except AFR have the reference allele and all super-populations have an alternative allele that is not present in the databases.

## Discussion

Our research provides a tool to mine whole genome sequence data to collectively evaluate the status of HERV-K provirus at known polymorphic and fixed sites in the human genome. The tool incorporates a statistical clustering algorithm to accommodate low depth sequence data and a visualization tool to explore the co-occurrence of polymorphic HERV-K in the global populations represented in the KGP data. There are numerous significant differences in the prevalence of individual and co-occurring polymorphic HERV-K among the five KGP super-populations. It is notable that individuals from EAS carry a lower total burden of HERV-K than other represented populations. These data provide a comprehensive framework of HERV-K genomic diversity to advance studies on potential roles for HERV-K in human disease, which have been alluring yet difficult to establish [19,20,22].

Tools developed to interrogate ERV insertional polymorphism typically exploit the unique signature created by the host-virus junction [11,44,45]. These approaches indicate that a site is occupied by an ERV but not whether there is a provirus associated with the site, which is more difficult to accomplish with short read sequence data. Our analysis tool provides an efficient means to detect occupancy and provirus status in one step. We decrease computational time by analyzing only the set of reads that map to existing HERV-K in the reference genome. This approach is justified because the polymorphic HERV-K that are missing from the human reference are closely related to those in the reference genome assembly and hence reads derived from them map to a related HERV-K in the reference. We employ *k-mer* counting methods, which also increase computational efficiency. A reference set of *k-mer*s that is unique to a HERV-K is generated for each location in the genome and the proportion of reads from the query set that maps to the *k-mer* reference set is reported as a continuous variable; there is no threshold of read count or depth imposed for classification. Instead we utilize a mixture model to cluster values and assign the same HERV-K status to the entire cluster. Clusters with n/T of 1 have all the unique *k-mer*s identified in the HERV-K reference set. We classify other clusters by determining if *k-mer*s mapped on the reference allele are distributed at sites in the coding portion of the genome or only in the LTR. This approach led to the interesting finding that several HERV-K could have population specific alleles.

Wildschutte *et al* [11] have conducted the most comprehensive study of HERV-K prevalence in the KGP data to date. While the goal of that paper was to identify new polymorphic insertions of either provirus or solo LTR in the KGP data, their analyses provide the prevalence of some polymorphic HERV-K provirus for comparison with our results. There are five HERV-K previously reported in Subramanian *et al* 2011 [10] that were not included in the Wildschutte paper [11]; all are polymorphic in our analysis (range 43-99%, see Table 1 and S1_Dataset:virus-column N). Seven polymorphic HERV-K, which Wildschutte *et al* [11] indicate occur in greater than 98% of KGP individuals, are fixed in our study. Our estimated prevalence for 14 HERV-K differs from that reported in Wildschutte et al [11] by 5% or more. Of these 14, the prevalence estimates at chr1:155596457-155605636 are most divergent. Our data show this site is fixed for provirus and Wildschutte *et al* [11] report that only 14% of the KGP data, all from AFR, have a HERV-K provirus integration. Our plots for chr1:155596457-155605636 show that AFR individuals carry the reference allele at this site (n/T near 1, Fig 5A) and all other individuals have n/T near 0.5. The *k-mer*s from individuals with low n/T values for chr1:155596457-155605636 map to only a subset of sites marked by unique *k-mer*s in the coding region (S8 Fig), which is consistent with sequence polymorphism or a deletion at these positions. The reference set T is small for this HERV-K and therefore overall coverage of the genome is low. Because Wildschutte *et al* [11] used a minimum coverage threshold for their *k-mer* mapping method, it is possible that alleles present in non-AFR populations would be outside their inclusion criteria. There is a similar signal for alleles, represented by lower n/T values, at the other 13 HERV-K sites although the differences between our prevalence estimates and those of Wildschutte et al [11] are small (S1_Dataset:virus). In most cases these putative alleles are found in all populations at different frequencies but in five there is some degree of population specificity (Fig 5, S7 Fig, S1_Dataset:virus). Our results indicate that there could be considerably more sequence variation in HERV-K among human populations than previously appreciated. These data also suggest that using HERV-K consensus sequences to study pathogenic potential could miss important features of HERV-K polymorphism, which can be characterized by both the site occupancy status (presence/absence) and, when present, by sequence differences in among individuals.

HERV-Ks are the youngest family of endogenous retroviruses in humans and consequently they share considerable sequence identity. This has the effect of limiting the number of unique sites associated with some HERV-K, which decreases the size of the reference set T (S1_Dataset:virus). Another example of a polymorphic HERV-K with a small set T is chr8:12316492, reported to be human specific (9), which shares a recent common ancestor with two older HERV-K (chr8:12073970 and chr8:8054700) all located at 8p23.1. Our data indicate that 14% of KGP individuals have the reference allele and most n/T values are less than 0.4 and fall into two non-zero clusters (S9 Fig). These appear to represent various structural variations (truncation or deletion) because there are several peaks in both the LTR and in the 5’ coding region. Thus although an n/T ratio of 0 or 1 reliably indicates absence and presence of the reference HERV-K, respectively, when T is small, sequence polymorphism and a deletion event can be difficult to distinguish from a solo LTR. However, because our mixture model statistically clusters similar n/T values, all individuals in a cluster have the same status (e.g allele or solo LTR) even if we do not know what that state is.

Our approach provides human disease researchers with a rapid means to determine if the frequency, and overall burden of the 96 HERV-K proviruses evaluated differ between a patient data set and populations represented in KGP. The visualization tool allows investigators to determine if HERV-Ks co-occur in certain clinical settings. The potential that HERV-K has multiple allelic forms in different populations is worthy of further analysis because a sequence allele could also contribute to a disease condition.

## Materials and methods

### HERV-K proviruses

The 96 HERV-K proviruses previously reported [10,11,32,46] were supplemented with HERV-K alleles present in the NCBI nt database (November 2016 release). We required that any allele of a HERV-K from the nt database have at least 2kb of reference-matching host flanking sequence to confirm genome location. In total, 234 alleles were collected at the 96 known HERV-K loci (92 in hg19, and 4 from the nt database). The location information and virus features are summarized in S1_Dataset: virus.

### Developing a *k-mer* based detection model

We identified the *k-mer*s that correspond to unique sequence characterizing each HERV-K. *K-mer*s are substrings (subsequences) of length *k* that exist in a string (DNA sequence). The length *k* is determined empirically (S1 Method). Each *k-mer* is labeled with the corresponding viruses in which it is observed.

Only those *k-mer*s referring to a single virus, unique *k-mer*s, are selected for the set T. Where multiple alleles of a HERV-K are available, *k-mer*s unique to all alleles at that location comprise T. Multiple 2bps different *k-mer*s (such as SNPs) corresponding to the same location on the virus, are merged into a single entry for the purposes of computing T. We map unique *k-mer*s back to the corresponding alleles to determine depth of the HERV-K (S3 Fig) and whether *k-mer*s are located in LTRs. (S1_Dataset: virus)

### Analysis of 1000 Genome Project (KGP) Data

To develop a method to recover sequences containing information on HERV-K we leverage the fact that HERV-Ks are closely related. Thus, most sequence reads obtained from an individual with a polymorphic HERV-K that is absent in the human reference, hg19, will map to the location of one of the closely related HERV-K that is present the human genome reference. A file with the coordinates for all reported HERV-K insertions is used to extract mapped reads from a genome sequence file (S1_Dataset:bed, which provides the coordinates for both hg19 and hg38). Note that the KGP data were mapped to GRCh37, which includes the decoy sequence hs37d5. This decoy contains the HERV-K at chr1:73594980_73595948 and is not present in hg19. Thus, we did not recover any reads for this HERV-K, which is polymorphic but reportedly at high prevalence in most populations [11].

The KGP data were downloaded in aligned Binary Alignment/Map (BAM) format (ftp://ftp.ncbi.nlm.nih.gov/1000genomes/ftp/data/). It contains data for 2,535 individuals (S1_Dataset:KGP) sequenced via low-depth whole-genome sequencing (mean depth = 6.98X). The individuals represent 26 populations, derived from 5 super-populations, including African (AFR), Admixed America (AMR), East Asian (EAS), European (EUR), and South Asian (SAS) [47,48]. Of 2,535 individuals, 28 also have high-depth DNA sequences (mean depth = 48.06X), which we use as a pilot dataset for our clustering methods, described below and in Supplementary Methods.

Our computational framework to indicate the status of each known HERV-K provirus is based on the *n/T* ratio, which is the proportion of *k-mer*s from each individual that are identical to the reference set T for each HERV-K. Sequence reads are extracted from a mapped file of whole human genome sequence data based on coordinates corresponding to each annotated HERV-K. The reads are *k-mer*ized and mapped to the set T, which represents all unique *k-mer*s assigned to each HERV-K in the reference set. We use exact match to map *k-mer*s data set to the unique *k-mer*s references. The *n/T* ratio is used as an indicator of the presence of each HERV-K. Using a hash table (S5 Method), it takes 15 minutes to generate the n/T matrix for 100 files. The source code for the entire process is at https://github.com/lwl1112/polymorphicHERV

### Dirichlet process Gaussian mixture model (DPGMM)

Because n (the number of *k-mer*s obtained from a persons’ sequence data, that map to a specific HERV-K) is affected by sequencing depth, we utilized a statistical model to cluster n/T for each HERV-K for each individual and assigned HERV-K status to a cluster. Mixture modeling is arguably the most widely used statistical method for clustering. In this analysis, we follow the work proposed by Lin et al. 2015 [49], which employs a Gaussian Mixture Model (GMM) with density function given by

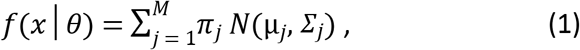

and with a mixture of relatively large number M Gaussian components (denoted by *N*(μ*j*, *Σj*), for j = 1:M) to represent the data density. To allow a flexible modeling approach, we employ the standard Bayesian (truncated) Dirichlet Process prior for the parameters *θ* = (*π_j_*,*μ_j_*, *Σ_j_*, *j* = 1:*M* [50,51]. The idea is that some of the mixture probabilities (*π_j_*) can be zero, hence the actual number of mixture components needed may be smaller than the upper bound M. This mechanism allows automatic determination of the number of mixture components needed by the data set at hand. Given a fitted model via the Bayesian expectation–maximization algorithm, in terms of estimates of all parameters *θ*, we identify clusters by aggregating Gaussian components. Merging components into clusters can be done by associating each of the Gaussian components to the closest mode of f(x|*θ*). Hence, the number of modes identified is the realized number of clusters. [See S2 Method for full detail]

### Co-occurrence of polymorphic HERV-K

We consider that both the individual frequency of a HERV-K and the co-occurrence of multiple HERV-K could differ among populations.

The time of a brute-force approach for finding all combinations *C_m_* of size m from p polymorphic HERV-K is 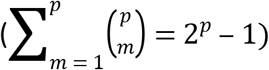, which is not efficient and redundant. We employed the Apriori algorithm [52], which is commonly used for finding frequent pattern sets; in our case indicating which polymorphic HERV-K frequently appear together. It first generates combinations C_m_ (initialized to 1). In the optimization, frequent combinations F_m_ are returned from candidates C_m_ when frequency exceeds the minimum threshold of co-occurrence. F_m_ are then self-joined to generate combinations C_m + 1_ of size *m* +1 and out of which F_m + 1_ satisfy the minimum co-occurrence. In each pass, candidate combinations are pruned so as to avoid generating all combinations, which reduces running time significantly.

### Statistical analysis of HERV-K frequencies across populations

We make statistical comparisons across 5 super-populations for the following three problems. For each problem, there are 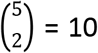 families of 1-to-1 comparisons conducted. The ‘prop-test’ function in R is used to test whether the proportions for two super-populations are the same.

1) individual prevalence of polymorphic HERV-K. (20 comparisons for each polymorphic HERV-K in a family)

2) the number of polymorphic HERV-K present per individual. (21 comparisons as the number of co-occurring polymorphic HERV-K is from 0 to 20)

3) the co-occurrence for combinations of polymorphic HERV-K.

Therefore, multiple hypotheses would be conducted on frequencies *F* across super-populations *P*_1…5_ as follows:

Null hypothesis, 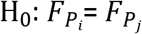, where i ≠ j;

Alternative hypothesis, 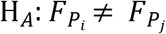, where i ≠ j.

A separate P-value is computed for each test and the Benjamini-Hochberg procedure [53] is used to account for multiple comparisons.

### Visualization in D3.js

We utilized D3.js (Data Driven Documents) [54], an open-source java script library to create an interactive visualization to display co-occurrence of polymorphic HERV-Ks in human populations. Our visualization system includes two modules, a welcome page and a result page. Input JSON data include locations of polymorphic HERV-K, population information, and the 0/1 (absence / presence) matrix. (See S3 Method). Source code is available at:

https://github.com/lwl1112/polymorphicHERV/tree/master/visualization

## Acknowledgements

WL was supported in part by the Louis S. and Sara S. Michael Endowed Graduate Fellowship in Engineering and the Fred A. and Susan Breidenbach Graduate Fellowship in Engineering.

